# Neuraminidase1 Activity Contributes to Vasopressin Receptor-mediated Augmentation of Water and Electrolyte Retention by the Kidney in *Eln* Haploinsufficient Mice

**DOI:** 10.64898/2026.06.11.731713

**Authors:** Gagandeep Kaur, Akua Serwaa-Bonsu, Kisho Miyasako, James A. McCormick, Patrick Osei-Owusu

**Affiliations:** Department of Physiology & Biophysics, Case Western Reserve University School of Medicine, Cleveland, Ohio, United States; Department of Medicine, Oregon Health & Science University, Portland, Oregon, United States

**Keywords:** elastin haploinsufficiency, vasopressin receptors, neuraminidase-1, renal function, extracellular fluid volume regulation

## Abstract

Elastin haploinsufficiency is a primary determinant of arteriopathy and hypertension that hallmark Williams syndrome (WS), a rare genetic disorder resulting from microdeletion of genes on human chromosome 7, including the elastin gene (*ELN*). Accumulating evidence suggests renal dysfunction, including enhanced sodium and water retention as an underlying cause of blood pressure elevation resulting from heterozygous deletion of *Eln* (*Eln^+/-^*) in mice that recapitulates the cardiovascular phenotype of WS. However, the underlying pathophysiological mechanisms are poorly understood. Here, we determined whether the activity of neuraminidase-1 (NEU1) of the elastin receptor complex (ERC) contributes to abnormal handling of water and electrolytes by the kidney in *Eln* haploinsufficiency. Adult male and female *Eln^+/+^* and *Eln^+/-^* mice were subjected to acute extracellular fluid volume expansion with normal saline, combined with pharmacological intervention targeting vasopressin V2 receptor (V2R), NEU1, ENaC, and NKCC2. In male *Eln^+/+^* mice, V2R blockade induced a dose-dependent increase in urine flow rate without affecting sodium excretion. Conversely, V2R stimulation with desmopressin markedly increased urinary sodium excretion in male *Eln^+/+^* but not *Eln^+/-^* mice, while both sexes of *Eln^+/-^* mice exhibited marked suppression of urine flow rate. Abrogation of ERC signaling through NEU1 inhibition produced a modest increase in urinary sodium excretion in male mice of both genotypes but augmented urine flow rate only in male *Eln^+/+^*mice. NEU1 blockade strikingly enhanced the natriuretic effect of furosemide and amiloride in male *Eln^+/+^* and modestly in *Eln^+/-^*mice. Taken together, we conclude that *Eln* haploinsufficiency disrupts vasopressin-dependent modulation of sodium and water reabsorption by sex-dependently altering ERC-mediated modulation of NKCC2 and ENaC. These findings reveal a novel mechanism by which abnormal ERC activity due to *Eln* haploinsufficiency potentially contributes to renal dysfunction and hypertension.

**GRAPHICAL ABSTRACT:** 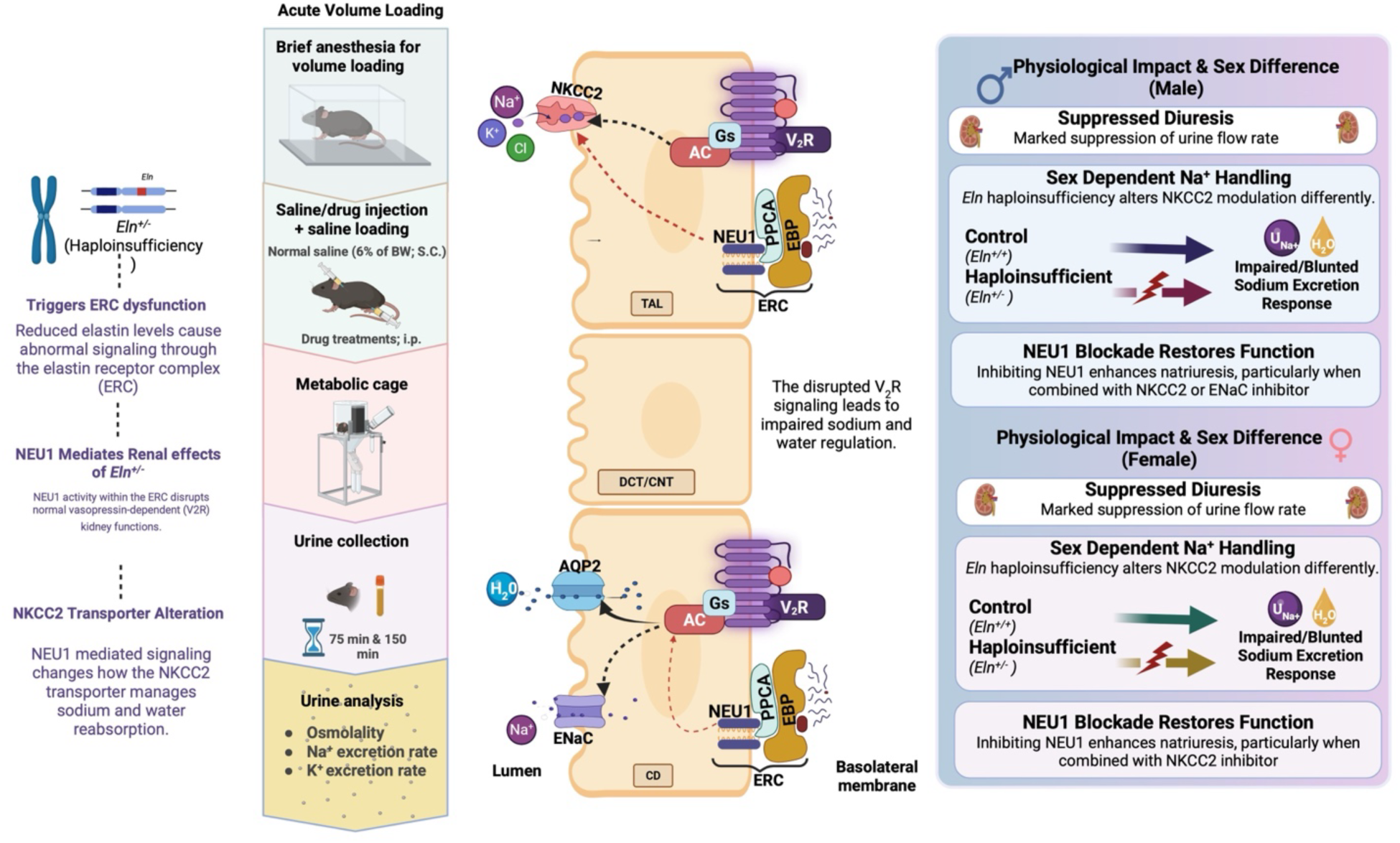

AC, adenylyl cyclase; AQP2, aquaporin 2; CD, collecting duct; CNT, connecting tubule; DCT, distal convoluted tubule; EBP, elastin binding protein; *Eln*, elastin allele; ENaC, epithelial sodium channel; ERC, elastin receptor complex; Gs, stimulatory Gα subunit; NEU1, neuroaminidase1; NKCC2, sodium-potassium-chloride cotransporter; PPCA, protective protein/ cathepsin A; TAL, loop of Henle thick ascending limb; V2R, vasopressin receptor type 2

## INTRODUCTION

The vasopressin system, activated by the anti-diuretic hormone, arginine vasopressin (AVP), plays a central role in maintaining fluid homeostasis through its regulation of renal water reabsorption and vascular tone. AVP is secreted in response to increased plasma osmolality and decreased plasma volume, and acts primarily on high-affinity vasopressin V2 receptors (V2R) located on the basolateral membrane of collecting duct principal cells and low-affinity V1R on vascular smooth muscle cells of the resistance vasculature^1-6^. V1R stimulation by AVP induces calcium release from intracellular stores leading to vasoconstriction and increased peripheral vascular resistance, in response to hypovolemia to maintain systemic blood pressure and visceral organ perfusion^3,4,7^. Activated V2R stimulates adenylyl cyclase leading to elevated intracellular cyclic adenosine monophosphate (cAMP) levels, which in turn activate protein kinase A (PKA)^8,9^. This cascade promotes the phosphorylation and apical membrane insertion of aquaporin-2 (AQP2) water channels, thereby enhancing water permeability and reducing urine volume while increasing urine osmolality^10,11^.

Besides water reabsorption, V2R signaling has been implicated in tubular sodium handling through the regulation of sodium transport via sodium-potassium-chloride cotransporter 2 (NKCC2) in the thick ascending limb and epithelial sodium channel (ENaC) in the connecting tubule and collecting duct^12-14^. Previously, it was reported that V2R activation promotes ENaC insertion in the apical membrane of principal epithelial cells, thus enhancing Na^+^ reabsorption^15^. Accordingly, constitutive activation of V2R due to genetic mutations or other factors can cause non-osmotic water reabsorption and plasma volume expansion, thus inducing Liddle syndrome-like hypertension^1,16^. Evidence from a previous study indicates that hypertension resulting from *Eln* haploinsufficiency (*Eln^+/-^*) in mice is associated with V2R-dependent enhanced retention of excess Na^+^ and water, suggesting a volume-dependent hypertension^17^.

Elastin is a key extracellular matrix (ECM) protein, which confers vascular compliance, most notably in the aorta^18-20^. Besides the structural role, elastin also serves as a source of biologically active molecules. Proteolytic processing of tropoelastin and mature elastin by extracellular matrix (ECM) proteases, including neutrophil elastase, cathepsins, and several matrix metalloproteinases, generates elastin-derived peptides (EDPs), many of which function as matrikines^21,22^. These biologically active EDPs, often containing conserved GxxPG or VGV motifs, elicit cellular responses primarily through activation of the broadly expressed heterotrimeric elastin receptor complex (ERC) located at the plasma membrane^23-25^. The ERC comprises elastin-binding protein (EBP) which mediates ligand recognition, protective protein/cathepsin A (PPCA), and neuraminidase-1 (NEU1) whose activation initiates downstream signaling cascades that regulate vascular reactivity, cell migration, proliferation, inflammation, and ECM remodeling^22,26-29^. ERC signaling has also been shown to be particularly relevant in conditions associated with increased elastin degradation such as aging, vascular disease, and cancer^22,23,30,31^. *Eln* haploinsufficiency, a condition characterized by reduced vascular elastin content, has been associated with accelerated cardiovascular and renal aging characterized by marked fragmentation of mature vascular elastin, increased arterial stiffness, and impaired renal hemodynamics^19,32,33^. Moreover, in a recent study, we reported that enhanced retention of excess Na^+^ and water in *Eln ^+/-^* mice is associated with blunted diuresis induced by the V2R antagonist, tolvaptan^17^. This suggests that *Eln* haploinsufficiency alters the regulation of renal tubular function by V2R; However, the mechanisms remain poorly understood. In this study, we examined how *Eln* haploinsufficiency affects renal handling of excess water and sodium upon V2R activation. We also assessed whether the putative signaling pathways that mediate the cellular effects of EDPs play any role in tubular sodium and potassium transport. Our results show that *Eln* haploinsufficiency heightens renal sensitivity to V2R activation and is accompanied by augmented sodium and potassium reabsorption via NKCC2 and ENaC. Moreover, we found that *Eln* haploinsufficiency promotes the reabsorption of excess water and sodium, and that this effect appears to be enhanced by signaling that mediates the physiological effects of EDPs.

## MATERIALS AND METHODS

Studies were performed in accordance with protocols approved by the Institutional Animal Care and Use Committee of Case Western Reserve University School of Medicine. Experiments were performed using 2–4-month-old male and female wild-type (*Eln^+/+^*) and *Eln* heterozygous (*Eln^+/-^*) mice that have been backcrossed more than ten generations into the C57BL/6 genetic background (Charles River). The generation of *Eln^+/-^* mice has been described previously^19,34^. Mice were provided unlimited access to food and water in our institution’s animal resource center at 22° C with a 12-h light/dark cycle.

### Measurement of urine parameters after expansion of extracellular fluid volume

Adult male and female *Eln^+/+^* and *Eln^+/-^*mice were administered various pharmacological agents via intraperitoneal (i. p.) injection. The agents included normal saline (vehicle control) at a volume of 100 µL (Cat No: S4041, Teknova; Hollister, CA, USA), tolvaptan (1, 2 and 3 mg/kg body weight [BW]; Cat. No. 5181, Tocris Biotechne, Minneapolis, MN, USA), desmopressin ([deamino-Cys^1^, D-Arg^8^]-Vasopressin acetate salt hydrate, dDAVP, 0.02, 0.06, and 0.2 µg/kg BW; Cat No. V1005-1MG, Sigma-Aldrich, St. Louis, MO, USA), amiloride hydrochloride (2 mg/kg BW; Cat No: 2016-88-8, Tocris Biotechne, Minneapolis, MN, USA), furosemide (10 mg/kg BW; Cat No: F4381-1G, Sigma-Aldrich, St. Louis, MO, USA), 2-deoxy-2,3-didehydro-N-acetylneuraminic acid (DANA,10mg/kg BW; Cat No: D9050-5MG, St. Louis, MO, USA), amiloride (2 mg/kg BW) + DANA (10 mg/kg BW), or furosemide (10 mg/kg BW) + DANA (10 mg/kg BW). All injections were prepared in sterile normal saline and administered in a total volume of 100 µL per mouse. Immediately following the i. p. injection, each mouse received a subcutaneous bolus injection of 5% dextrose in normal saline; The volume of this injection was calculated as 6% of the animal’s body weight to ensure consistent fluid loading across subjects. This procedure was performed under light isoflurane anesthesia to minimize stress and discomfort during administration. After treatment, mice were promptly transferred to individual metabolic cages designed to allow precise urine collection, without access to food or water to prevent confounding variables related to intake. Urine samples were collected at two time points: 75 minutes and 150 minutes post-treatment and acute volume loading. Upon completion of the urine collection period, all animals were returned to their respective home cages and monitored for recovery. Urine osmolality was measured immediately using VAPRO® Vapor Pressure Osmometer.

### Measurement of urine sodium and potassium excretion rate after acute extracellular fluid volume expansion

After collection in metabolic cages, total urine volume was recorded, and samples were centrifuged at 3,000 rpm for 10 minutes to remove debris. Sodium and potassium concentrations were measured using flame photometry, as previously described^16,17^, with the user blinded to the sample identity. Samples were diluted as needed to ensure accurate readings within the instrument’s linear range. Urine excretion rates of sodium (U_Na_^+^) and potassium (U_K_ ^+^) were normalized to BW in grams.

### Glomerular filtration rate measurement in conscious mice

Glomerular filtration rate (GFR) was measured using the transdermal FITC-sinistrin clearance method by following the protocol described by Scarfe et al., 2018^35^. Briefly, FITC-sinistrin solution (40 mg/mL) was prepared in normal saline and protected from light. The mice were anesthetized with isoflurane for dorsal hair shaving followed by brief depilatory treatment one day before GFR measurement. The following day, a transdermal GFR monitoring device was affixed to the depilated flank, ensuring proper LED alignment and secure but non-restrictive placement. Following a 3-minute baseline period, FITC-sinistrin (0.15 mg/g BW) was administered via a retro-orbital injection under isoflurane anesthesia, and transdermal fluorescence was recorded for 1.5 – 2 hours. Individual mice were placed in separate cages with additional enrichment and a few food pellets for the duration of the transdermal fluorescence recording. Devices were removed under brief anaesthesia and the acquired data was used to calculate GFR using the manufacturer-provided software (MB_Lab2, Medibeacon). GFR values were normalized to BW.

### Statistical analysis

All data are mean ± standard error of the mean (SEM). For the assessment of within-group effects of various drug treatments on urine flow rate, urine osmolality, GFR, U_Na_^+^, and U_K_ ^+^, two-way analysis of variance mixed-effect model was used, followed by Sidak *post hoc* tests. The level of significance was set at *P* < 0.05.

## RESULTS

### *Eln* haploinsufficiency augments vasopressin receptor-dependent retention of excess water and sodium following acute expansion of extracellular fluid volume

The vasopressin system is recognized a major contributor to the regulation of extracellular fluid volume (ECFV) due to water and sodium reabsorption in the distal nephron^2,9,11,12^. Previously, we reported that *Eln* haploinsufficiency blunted the diuretic effect of V2R blockade with tolvaptan, accompanied by accentuated epithelial sodium channel (ENaC)-dependent sodium and water reabsorption in female mice on low-sodium diet^17^. These observations led to the conclusion that altered sodium and water homeostasis contributes to blood pressure elevation in *Eln^+/-^* mice^17^. Vasopressin has been shown to affect sodium reabsorption by upregulating ENaC activity in the principal cells of the connecting tubule (CNT) and collecting duct (CD)^15,36,37^. Indeed, certain mutations in V2R that lead to increased ENaC activity are implicated in volume-dependent hypertension^16,38^. V2R signaling is a key regulator of renal water reabsorption and have also been reported to indirectly influence ENaC activity^12,39^. Therefore, we determined whether *Eln* haploinsufficiency promotes ECFV expansion by altering the effects of vasopressin on renal sodium and water handling. We examined urine flow rate, osmolality, and urine sodium and potassium excretion rates following a bolus subcutaneous administration of normal saline administration (6% of body weight, SQ), in the presence or absence of the V2R agonist, dDAVP (0.02, 0.06 and 0.2 µg/kg, i.p.). In female mice, there was a significant effect of dDAVP injection (*P* = 0.0012) but not genotype (*P* = 0.8753) on urine flow rate, and no significant interaction between treatment and genotype (*P* = 0.3247). The lowest dose of dDAVP (0.02 µg/kg) led to a significant decline in urine flow rate (*P* = 0.0093), which remained markedly low at subsequent doses in *Eln^+/+^*mice (Figure 1A). Although *Eln^+/-^* mice demonstrated a similar trend towards reduced urine flow rate, the decrease was not significant (*P* = 0.9155) (Figure 1A). Female *Eln^+/+^*mice exhibited a rapid and sustained elevation in urine osmolality starting at the lowest dose, indicative of enhanced urine water reabsorption in response to V2R stimulation by dDAVP. By contrast, female *Eln^+/-^*mice showed only a modest osmolality response, suggesting a blunted or dysregulated osmoregulatory mechanism (Figure 1B). Furthermore, there was a significant effect of dDAVP (*P* = 0.0247) and genotype (*P* = 0.0018), as well as a significant interaction between dDAVP treatment and genotype (*P* = 0.0289) on urinary sodium excretion in female mice. In *Eln^+/+^* mice, there was a dose-dependent increase in urinary sodium excretion, whereas in *Eln^+/-^* mice, there was a substantial blunting of the natriuretic response to acute ECFV expansion across all doses of dDAVP, when compared to the response in *Eln^+/+^*mice (Figure 1C). Similarly, urine potassium excretion rate increased significantly in *Eln^+/+^*mice. Notably at 0.06 µg/kg dDAVP, urinary potassium excretion was significantly reduced compared to baseline in this genotype. In contrast, *Eln^+/-^* mice showed minimal changes across all doses of dDAVP, with a consistently lower urine potassium excretion rate relative to *Eln^+/+^* mice, highlighting a genotype-dependent impairment in renal potassium handling (Figure 1D). Overall, these findings indicate that V2R activation unmasks functional differences in renal sodium and potassium handling due to *Eln* haploinsufficiency in female mice, with *Eln^+/-^* mice showing impaired responses relative to *Eln^+/+^*mice.

**Figure 1.**
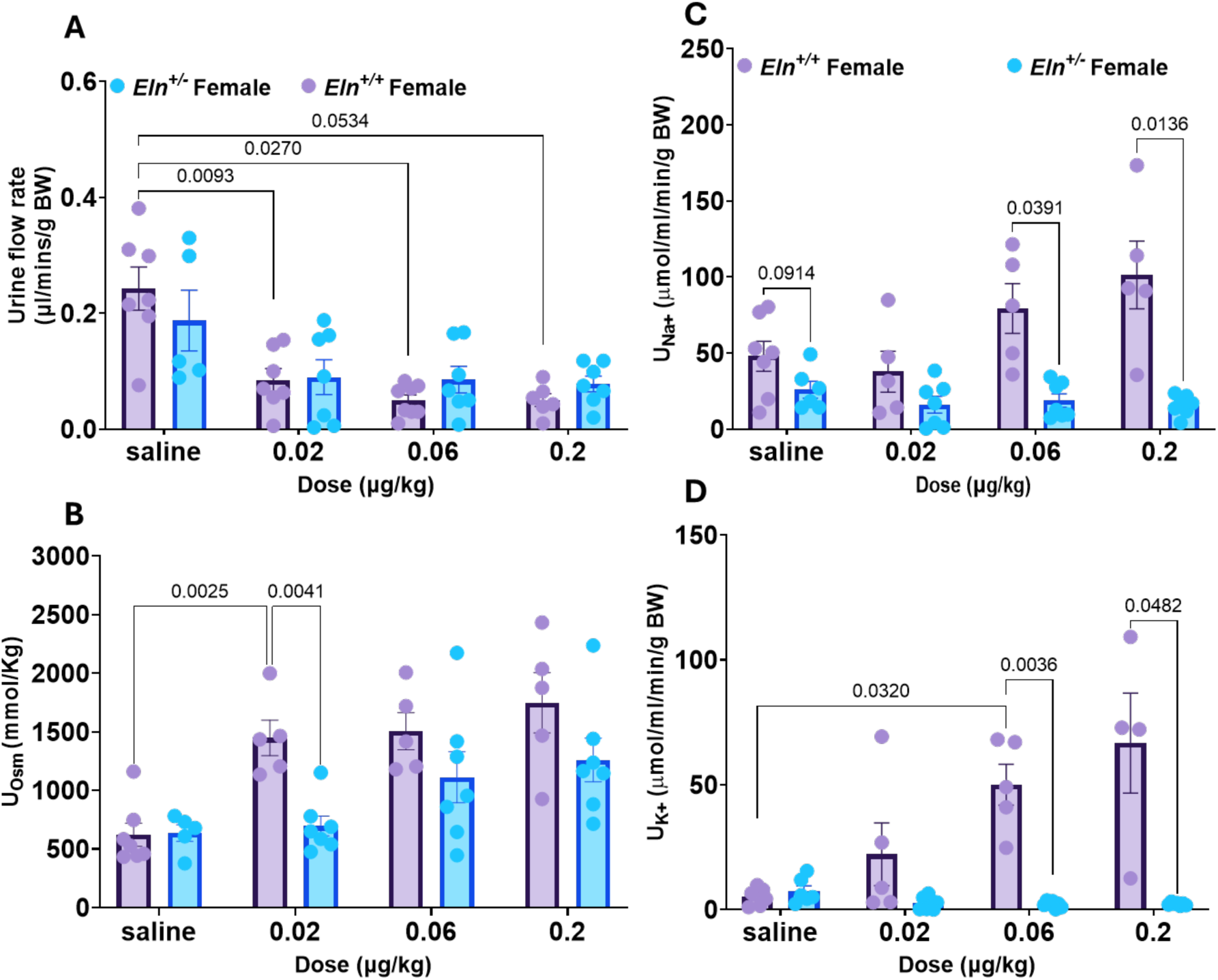
Urine flow rate, osmolality, and sodium and potassium excretion rate following acute extracellular fluid volume expansion with normal saline in the presence and absence of dDAVP (0.02, 0.06 and 0.2 µg/kg, i.p.) in female mice. (**A**) Urine flow rate after volume loading with saline, both with and without dDAVP, are presented for *Eln*^+/+^ and *Eln*^+/-^ mice. (**B)** Bar graph showing urine osmolality (U_osm_). (**C & D**) Bar graph showing urine sodium (U_Na+_) and potassium (U_K+_) excretion rate, respectively, which was calculated using the concentration of electrolytes in urine samples collected from the 75^th^ and 150^th^ minute. Symbols represent individual animals in each experimental group and genotype. Data were analyzed using two-way ANOVA mixed model with Sîdak post hoc test. Values in bar graphs are expressed as mean ± SEM.

As in female mice, we also determined how *Eln* haploinsufficiency affected diuresis and natriuresis following acute ECFV expansion in the presence of V2R agonist in male mice. Both dDAVP and genotype had a significant effect on urine flow rate (*P*<0.0001 and *P* = 0.0363, respectively), with a significant interaction between the two factors (*P* = 0.0064). In *Eln^+/+^* mice, urine flow rate declined significantly in a dose-dependent manner compared to baseline, whereas *Eln^+/-^* mice exhibited an exaggerated antidiuretic response at lower dDAVP doses (0.02 and 0.06 µg/kg), compared to *Eln^+/+^* males, indicating that *Eln* haploinsufficiency enhances the basal antidiuretic effect of V2R activation (Figure 2A). Both genotypes showed increased urine osmolality at all doses relative to saline controls, reaching significance at the highest dDAVP dose, i.e. 0.2 µg/kg (Figure 2B). Furthermore, there was a robust effect of genotype (*P* < 0.0001) but not dDAVP (*P* = 0.1320) and a significant dose-genotype interaction (*P* = 0.0378) on urinary sodium excretion, indicating altered natriuretic regulation in male *Eln^+/-^* mice. dDAVP treatment led to increased urine sodium excretion rate in *Eln^+/+^* mice, whereas *Eln^+/-^* male mice exhibited markedly attenuated natriuresis with a significant decline at a higher dose (0.2 µg/kg) compared to their *Eln^+/+^* counterparts (Figure 2C). For potassium excretion, *Eln^+/+^*mice demonstrated significant increases following dDAVP treatment, especially at higher doses. In contrast, *Eln^+/-^* mice showed minimal changes across all doses, with values significantly lower than those in *Eln^+/+^* mice (Figure 2D). These findings reveal enhanced dDAVP-induced anti-diuretic effect due to *Eln* haploinsufficiency in male mice. The results also suggest that *Eln* haploinsufficiency impairs vasopressin-mediated sodium and potassium handling by the kidney.

**Figure 2.**
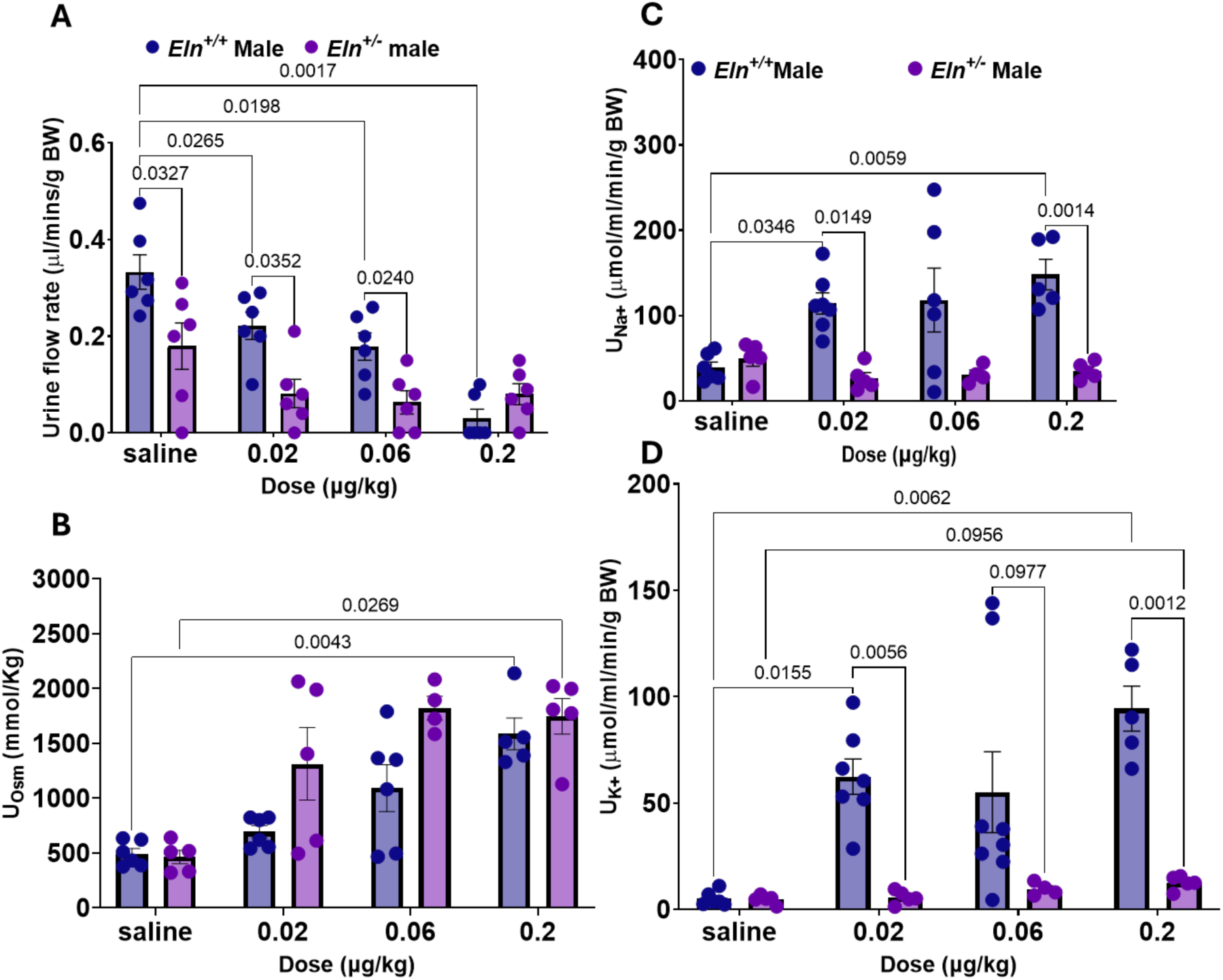
Urine flow rate, osmolality, and sodium and potassium excretion rate following acute extracellular fluid volume expansion with normal saline in the presence and absence of dDAVP (0.02, 0.06 and 0.2 µg/kg, i.p.) in male mice. (**A**) Urine flow rate after volume loading with saline, both with and without dDAVP, are presented for *Eln*^+/+^ and *Eln*^+/-^ mice. (**B)** Bar graph showing urine osmolality (U_osm_). (**C & D**) Bar graph showing urine sodium (U_Na+_) and potassium (U_K+_) excretion rate, respectively, which was calculated using the concentration of electrolytes in urine samples collected from the 75^th^ and 150^th^ minute. Symbols represent individual animals in each experimental group and genotype. Data were analyzed using two-way ANOVA mixed model with Sîdak post hoc test. Values in bar graphs are expressed as mean ± SEM.

### Sex-dependent impairment of NKCC2- and ENaC-mediated fluid and electrolyte regulation in *Eln* haploinsufficient mice

To further delineate the mechanism by which *Eln* haploinsufficiency leads to abnormal renal tubular function encompassing enhanced sodium and potassium reabsorption, we used in vivo pharmacology to interrogate ENaC and NKCC2 activity under acute volume loading. Mice were administered amiloride (2 mg/kg, i.p.) or furosemide (10 mg/kg, i.p.) to block ENaC at the distal nephron (CNT and CD) or NKCC2 in the thick ascending limb (TAL), respectively, immediately before acute subcutaneous volume loading with normal saline. In female mice of both genotypes, there was a significant effect (*P* = 0.0414) of furosemide and amiloride administration on urine sodium excretion rate and increasing trend when compared to excretion rate after administration of vehicle/saline (Figure 3A). For urinary potassium excretion, there was a significant effect of genotype in female mice (*P* = 0.0414) and a significant increase in potassium excretion rate in female *Eln^+/-^* mice after amiloride administration (Figure 3B). On the other hand, genotype had a robust effect (*P*<0.0001) on urinary sodium and potassium excretion in male mice of both genotypes. Both furosemide and amiloride elicited a pronounced natriuretic response in male *Eln^+/+^* and *Eln^+/-^* mice (Figure 3C). A similar response pattern was observed for urinary potassium excretion (Figure 3D). Collectively, these findings indicate sex-related differences in the sensitivity of *Eln* haploinsufficient mice to the natriuretic effects of NKCC2 and ENaC blockade.

**Figure 3.**
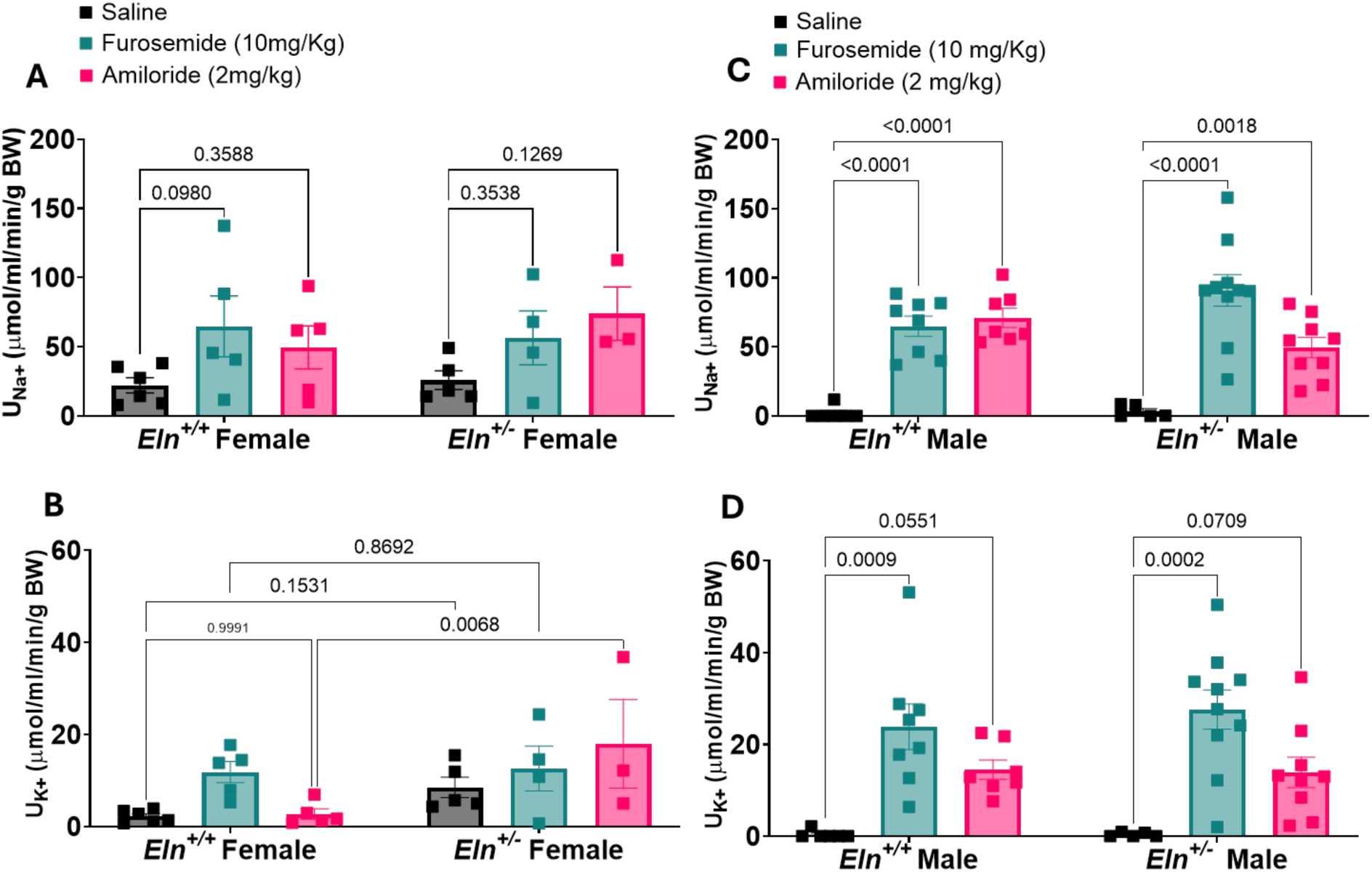
Urine sodium and potassium excretion rate following acute extracellular fluid volume expansion with normal saline with or without intraperitoneal injection of saline, furosemide, or amiloride in female and male *Eln*^+/+^ and *Eln*^+/-^ mice. (**A**) Urine sodium excretion rate (U_Na+_) is presented for *Eln*^+/+^ and *Eln*^+/-^ female mice. (**B)** Bar graph showing urine potassium excretion rate (U_K+_) in *Eln*^+/+^ and *Eln*^+/-^ female mice. (**C & D**) Bar graph showing urine sodium and potassium excretion rate in male *Eln*^+/+^ and *Eln*^+/-^ mice, respectively. Symbols represent individual animals in each experimental group and genotype. Values were calculated using the concentration of electrolytes in samples collected up to the 75^th^ minute. Data were analyzed using two-way ANOVA mixed model with Sîdak post hoc test. Values in bar graphs are expressed as mean ± SEM.

### *Eln* haploinsufficiency abolishes the diuretic effects of NEU1 blockade

Enzyme-mediated degradation of extracellular matrix (ECM) proteins such as collagen, mature elastin, and tropoelastin leads to the generation of matrikines, including biologically active elastin-derived peptides (EDPs) that elicit their physiologic effects via the broadly expressed heterotrimeric elastin receptor complex (ERC), comprising elastin-binding protein (EBP), protective protein/cathepsin A (PPCA), and neuraminidase-1 (NEU1)^22,26-28^. As previously reported, *Eln* haploinsufficiency is associated with a high degree of vascular elastic fiber fragmentation, suggesting increased rate of elastin degradation that likely alters the steady-state generation and effects of EDPs^19,22,34,40,41^. To determine whether altered EDP-ERC signaling plays a role in renal sodium and water handling abnormalities resulting from *Eln* haploinsufficiency, we interrogated the ERC signaling pathway in vivo using the NEU1 antagonist, 2,3-dehydro-2-deoxy-N-acetylneuraminic acid (DANA), in the acute volume-loading protocol that was used to probe vasopressin-dependent mechanisms. Additionally, we examined how targeted blockade of segment-specific sodium transport influenced electrolyte handling in *Eln^+/+^* and *Eln^+/-^*mice, with the goal of defining the contribution of altered EDP levels to renal sodium and potassium mishandling due to *Eln* haploinsufficiency. Mice received DANA (10 mg/kg, i.p.) alone or in combination with agents that inhibit sodium reabsorption in discrete nephron segments – amiloride (2 mg/kg, i.p.) to block ENaC in the CNT and CD, and furosemide (10 mg/kg, i.p.) to block NKCC2 in the TAL – followed immediately by acute subcutaneous volume loading as described above. In female mice, there was a significant effect of treatment (*P* = 0.0066) but not genotype (*P* = 0.2692) and no genotype–treatment interaction (*P* = 0.503). From post-hoc analysis, DANA administration alone or co-administration with amiloride did not increase urinary sodium excretion in either *Eln^+/+^* or *Eln^+/-^*mice (Figure 4A). However, following furosemide and DANA co-administration, there was a significant increase and an increasing trend in urine sodium excretion rate in *Eln^+/+^* and *Eln^+/-^* mice, respectively, compared to groups receiving saline/control or DANA alone (Figure 4A). For urinary potassium excretion rate, furosemide alone produced a rise, whereas amiloride had no effect in *Eln^+/+^* mice. In contrast, furosemide had no effect on urine potassium excretion rate in *Eln^+/-^*mice (Figure 4B). In female *Eln^+/+^* mice, DANA enhanced the natriuretic and kaliuretic effects of furosemide (Figure 3A & 4A), suggesting that signaling downstream of NEU1 promotes sodium and potassium reabsorption via NKCC2 in the TAL. In male mice, there was a significant effect of treatment (*P* < 0.0001) and genotype (*P* = 0.064), and a significant interaction between genotype and treatment (*P* = 0.0459). From post-hoc analysis, DANA alone caused a mild increase in urinary sodium excretion in both genotypes (Figure 4C). Co-administration of DANA and amiloride did not alter urinary sodium excretion in either genotype (Figure 3C vs 4C). In contrast, co-administration of DANA and furosemide produced a synergistic natriuretic effect in *Eln^+/+^* mice, and a strong but less robust effect in *Eln^+/-^* mice (Figure 3C & 4C). In *Eln^+/+^* mice, urinary potassium excretion rate increased significantly after DANA administration alone and after a combined administration of DANA and furosemide, with a similar but attenuated response in *Eln^+/-^*male mice (Figure 4D). Collectively, these results support the hypothesis that ERC signaling intersects with ENaC- and NKCC2-mediated transport and further demonstrate that the renal tubular consequences of ERC inhibition are affected by sex and *Eln* genotype, underscoring a complex regulatory interplay between ERC activity and downstream sodium-handling pathways.

**Figure 4.**
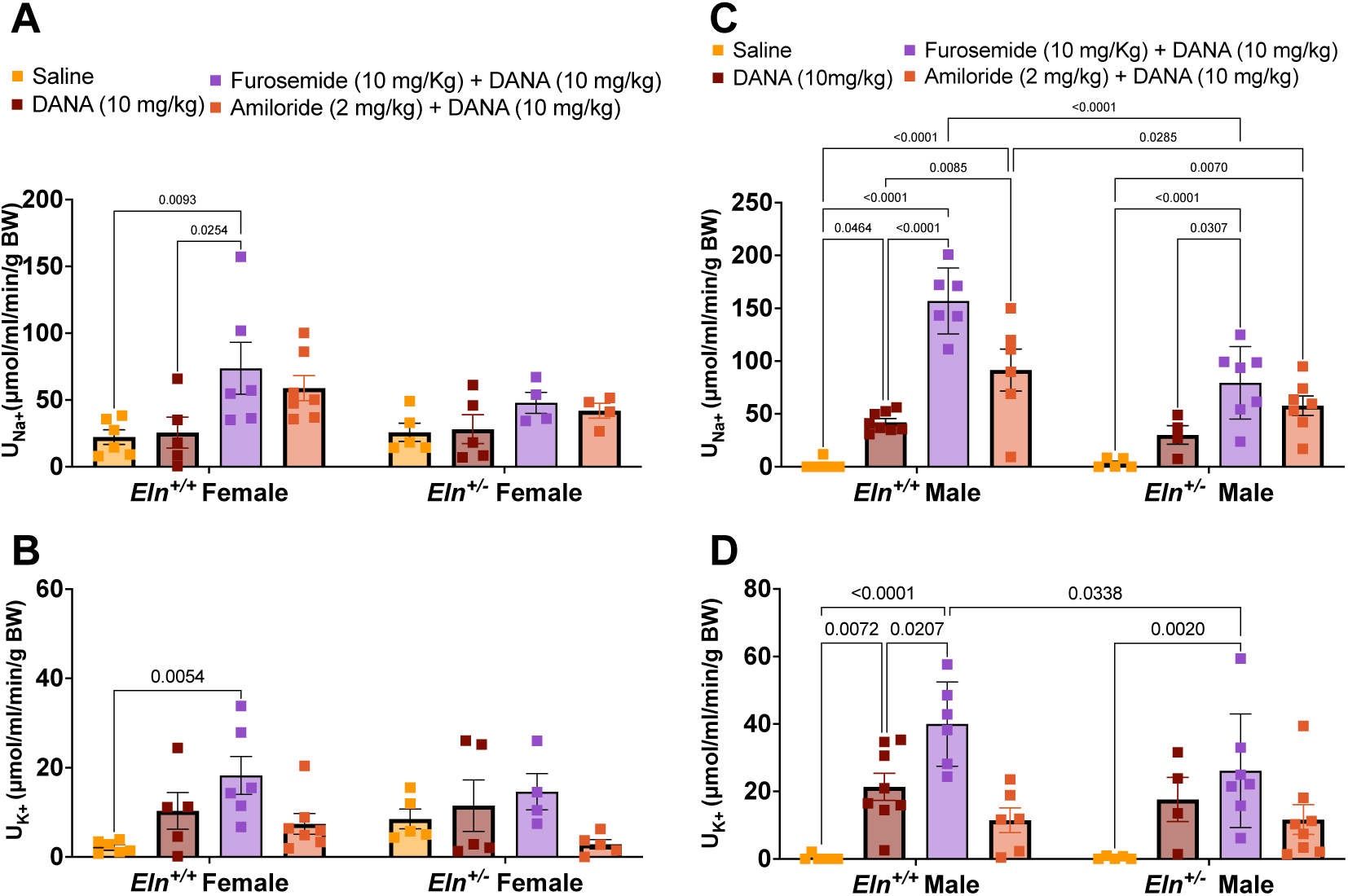
Urine sodium and potassium excretion rate following acute extracellular fluid volume expansion with or without intraperitoneal injection of furosemide, amiloride, furosemide + DANA, or amiloride + DANA in female and male *Eln*^+/+^ and *Eln*^+/-^ mice. (**A, C**) Bar graph showing urine sodium excretion rate (U_Na+_) in *Eln*^+/+^ and *Eln*^+/-^ mice. (**B, D)** Bar graph showing urine potassium excretion rate (U_K+_) in *Eln*^+/+^ and *Eln*^+/-^ mice. Symbols represent individual animals in each experimental group and genotype. Values were calculated using the concentration of electrolytes in urine samples collected from the 150^th^ minute. Data were analyzed using two-way ANOVA mixed model with Sîdak post hoc test. Values in bar graphs are expressed as mean ± SEM.

### NEU1 inhibition modulates the effect of V2R stimulation on urine sodium and potassium excretion rates in a sex- and genotype-dependent manner

The observations made so far show that enhanced sodium and water retention due to *Eln* haploinsufficiency is associated with altered vasopressin-mediated water and sodium handling and potentiation of sodium and potassium reabsorption via NKCC2 by ERC signaling. These results led us to ask whether ERC signaling intersects with vasopressin-evoked mechanisms to regulate sodium and water reabsorption and whether such an interaction is affected by *Eln* haploinsufficiency. Accordingly, we postulated that ERC activity enhances V2R-mediated regulation of renal sodium and water transport such that blockade of ERC signaling would attenuate vasopressin-evoked anti-diuretic and anti-natriuretic responses in *Eln^+/-^* mice. To test this hypothesis, we co- administered the NEU1 antagonist, DANA, and dDAVP and assessed the diuretic and natriuretic outcomes following acute ECFV expansion with normal saline. In female mice, 2-way ANOVA revealed a significant effect of treatment on urinary sodium excretion (*P* = 0.0162). There was no difference in urine sodium excretion rate between dDAVP alone and dDAVP + DANA in *Eln^+/+^* mice. In contrast, co-administration of dDAVP and DANA resulted in a robust increase in urine sodium excretion rate in *Eln^+/-^*mice, compared with saline/control or dDAVP alone (Figure 5A). For urine potassium excretion rate, co-administration of dDAVP and DANA led to a significant increase in *Eln^+/+^* but not *Eln^+/-^* mice (Figure 5B). Similar patterns in urine sodium and potassium excretion rates were observed in male mice, where there were a significant effect of treatment (*P* = 0.0047) and a strong genotype-by-treatment interaction (*P* < 0.0001). dDAVP alone markedly increased urine sodium excretion rate in *Eln^+/+^*mice, whereas this response was blunted in *Eln^+/-^* mice; However, co-administration of dDAVP and DANA markedly increased urinary sodium excretion in *Eln^+/-^* relative to *Eln^+/+^* mice (Figure 5C). Additionally, urinary potassium excretion increased significantly in both genotypes after dDAVP + DANA administration compared to saline (Figure 5D). Together, these findings support the conclusion that ERC signaling contributes to altered responsiveness to vasopressin in *Eln* haploinsufficiency, and that pharmacological inhibition of NEU-1 partially restores urinary sodium and potassium excretion in a sex- and genotype-dependent manner.

**Figure 5.**
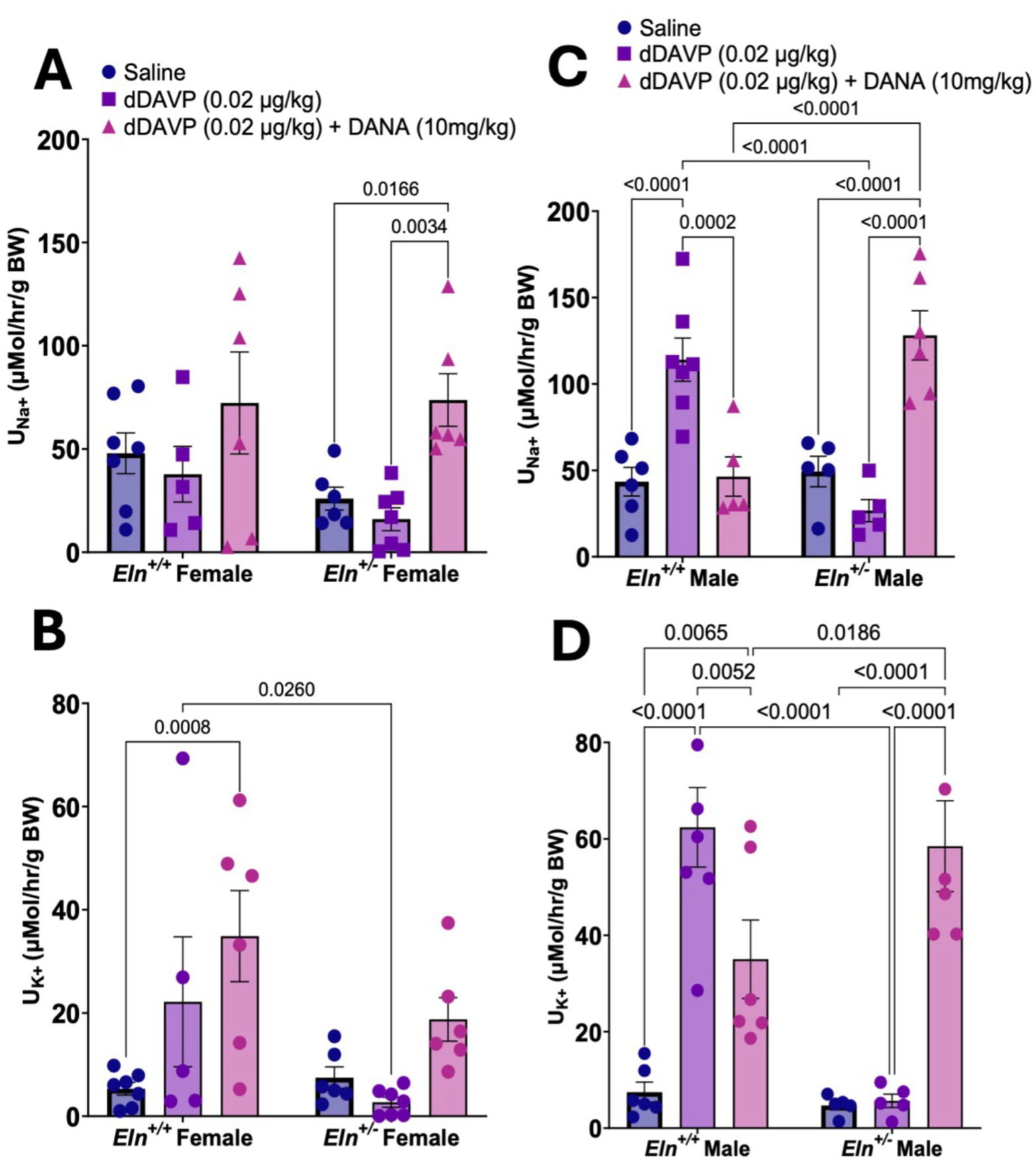
Urine sodium and potassium excretion rate following acute extracellular fluid volume expansion with or without intraperitoneal injection of dDAVP and dDAVP + DANA in female male *Eln*^+/+^ and *Eln*^+/-^ mice. (**A, B**) Bar graphs showing urine sodium (U_Na+_) and potassium (U_K+_) excretion rates in female *Eln*^+/+^ and *Eln*^+/-^ mice. (**C & D**) Bar graphs showing urine sodium and potassium excretion rate in male *Eln*^+/+^ and *Eln*^+/-^ mice. Symbols represent individual animals in each experimental group and genotype. Values were calculated using the concentration of electrolytes in combined urine samples collected from the 75^th^, 150^th^ and 225^th^ minute. Data were analyzed using two-way ANOVA mixed model with Sîdak post hoc test. Values in bar graphs are expressed as mean ± SEM.

### ERC blockade unmasks genotype**-** and sex**-**specific effects of *Eln* haploinsufficiency on glomerular filtration rate

Because ERC activation by EDPs has been shown to promote vasodilation^42,43^, we investigated whether the tubular effects of NEU1 blockade was due, at least partly, to a tubular response because of upstream changes in renal hemodynamics and/or GFR. To this end, we measured GFR before and after NEU1 blockade with i.p. DANA administration. In female mice, there was no significant effect of treatment or genotype on GFR (Figure 6A). In contrast, male mice exhibited a distinct pattern: there was a significant effect of treatment (*P* = 0.0272) and genotype (*P* = 0.0453) on GFR. Notably, *Eln^+/-^* mice showed increasing trends toward significant increase in GFR following DANA administration compared to their respective baseline values and to DANA-treated *Eln^+/+^* mice (Figure 6B). These findings suggest that ERC signaling likely contributes to baseline glomerular filtration dynamics in a sex- and genotype-dependent manner.

**Figure 6.**
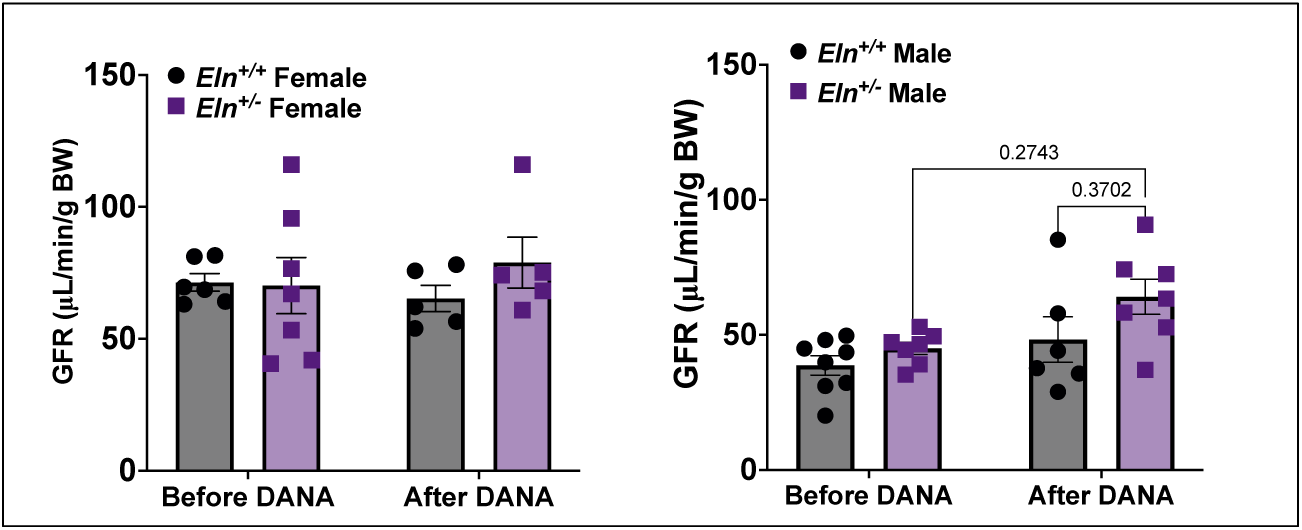
Transdermal glomerular filtration rate (GFR) before and after intraperitoneal injection of DANA in female and male *Eln*^+/+^ and *Eln*^+/-^ mice. (**A, B)** Bar graphs showing transdermal GFR in female and male *Eln*^+/+^ and *Eln*^+/-^ mice, respectively. Symbols represent individual animals in each experimental group and genotype. Data were analyzed using two-way ANOVA mixed model with with Sîdak post hoc test. Values in bar graphs are expressed as mean ± SEM.

## DISCUSSION

Hypertension is a hallmark of Williams syndrome (WS), a rare genetic disorder with characteristic ECM remodeling involving the fragmentation of vascular elastic fibers and proteolysis of ECM proteins including elastin (ELN). The blood pressure and vascular phenotype of WS mimics accelerated cardiovascular aging and age-related co-morbidities including isolated systolic hypertension and declining kidney function^19,33,34,40^. Kidney dysfunction associated with age-related hypertension is characterized by progressive loss of functional nephrons with a continuous decline in GFR, ultimately resulting in Na^+^ retention, ECFV expansion, and increased cardiac output that in turn increases or sustains resting blood pressure elevation in a vicious cycle^44-47^. Previously, we showed that hypertension in *Eln^+/-^* mice, the animal model of *Eln* haploinsufficiency, is sensitive to ENaC blockade and is also associated with blunted diuretic response to the blockade of vasopressin V2R with tolvaptan, suggesting excessive retention of sodium and water by the kidney^17^. Using an in vivo model of acute ECFV expansion, combined with renal tubular pharmacology, we show herein that increased sensitivity to V2R activation and renal tubular transport of sodium and potassium via NKCC2 and ENaC, sex dependently, mediates excessive retention of sodium and water by the kidney, in the context of *Eln* haploinsufficiency. Our results also show that signaling via the elastin receptor complex (ERC) likely contributes to the augmented V2R-mediated retention of sodium and potassium by the kidney in *Eln* haploinsufficiency.

Renal vasopressin signaling in principal cells of the distal nephron, mainly the collecting duct, is an established system for water conservation, and thus ECFV regulation by the kidney^10,11^. Classically, activation of basolateral V2R by vasopressin or its synthetic analog, desmopressin (dDAVP), activates the trafficking and apical membrane insertion of aquaporin 2 (AQP2) as water channels in CD epithelial principal cells, whereby the hypertonicity of the renal medullary interstitium provides the driving force for water reabsorption^10,11^. However, under conditions of genetic mutations or other factors that lead to constitutive V2R activation, ECFV expansion can also occur due to non-osmotic water reabsorption^38,48^. Interestingly, accumulating evidence directly implicates V2R activation in electrolyte transport in the loop of Henle TAL and the distal nephron segments including the CNT and CD^12,13,15,49^. Previous studies showed that V2R activation also promotes ENaC insertion in the apical membrane of CNT epithelial principal cells, thus enhancing Na^+^ reabsorption and inducing Liddle syndrome-like hypertension^8,50^. Vasopressin has also been shown to promote Na^+^ retention by stimulating NKCC2 to facilitate Na^+^ reabsorption in the TAL^12,49,51,52^. These previous studies demonstrate how the vasopressin system can co-opt NKCC2- and ENaC to promote Na^+^ reabsorption. The results in this study provide new evidence that *Eln* haploinsufficiency impinges on vasopressin-dependent water reabsorption mechanisms to positively regulate sodium and potassium reabsorption via NKCC2 and ENaC. Interestingly, V2R activation with desmopressin was paradoxically associated with a dose-dependent increase in urinary sodium and potassium excretion in *Eln^+/+^* mice upon acute ECFV expansion with normal saline. This may likely be due, partly, to a) rapid delivery of sodium and potassium to the distal nephron, including the TAL, distal convoluted tubule (DCT), CNT and CD, in amounts and rates exceeding the functional reabsorption limit of NKCC2, NCC, and ENaC in the respective nephron segments, and/or b) the anti-diuretic and urine concentrating effect of desmopressin, as revealed by the dose-dependent increase in urine osmolality. Interestingly, urinary excretion of both sodium and potassium was markedly attenuated in *Eln^+/-^*mice, despite the preserved anti-diuretic effect of desmopressin, which was enhanced in male animals. These results indicate that *Eln* haploinsufficiency enhances functional tubular sodium and potassium reabsorptive capacity. The results are also consistent with our prior study showing that *Eln*^+/-^ mice are insensitive to the diuretic effect of V2R blockade with tolvaptan^17^. Taken together, these lines of evidence suggest that *Eln* haploinsufficiency somehow alters vasopressin-mediated regulation of tubular water, sodium, and potassium handling.

Pharmacological evidence to support the mechanisms by which *Eln* haploinsufficiency enhances renal tubular sodium and potassium reabsorption included the elevation of urinary sodium and potassium excretion upon blockade of NKCC2 with furosemide and ENaC with amiloride. We observed a sex-related difference in the effect of NKCC2 and ENaC blockade on urinary sodium and potassium excretion, which were more robust in male relative to female mice of both genotypes. In addition, male *Eln^+/-^* mice were more responsive to furosemide and amiloride, suggesting an elevated NKCC2 activity due to *Eln* haploinsufficiency. The mechanism underlying the potential sex-related difference in NKCC2 and ENaC activity was not investigated in this study. However, the observation is consistent with the previously reported sex-related difference in the magnitude of systolic blood pressure elevation resulting from *Eln* haploinsufficiency in mice, which was attributed at least partly to blood pressure modulation by ovarian hormones^17,41^.

The effect of *Eln* haploinsufficiency on blood pressure and kidney function has been attributed largely to arteriopathy of conduit and resistance arteries associated with abnormal ECM remodeling, augmented vascular tone, and the vascular consequences of hyperactive renin-angiotensin system^34,41,53^. However, the ELN monomer, tropoelastin, and bioactive peptides derived from the degradation of mature, crosslinked ELN (elastin-derived peptides, EDPs) can elicit biological effects by activating plasma membrane-localized, widely expressed heterotrimeric ELN receptor complex (ERC)^22-25^. Thus, *Eln* haploinsufficiency likely alters ERC signaling due to changes in the steady-state level of bioactive EDPs. However, the role of ERC signaling in water and electrolyte regulation by the kidney is unclear. Here, blockade of the ERC signal transducing subunit NEU1 with DANA tended to increase GFR only in *Eln^+/-^*mice, which could result from vascular effects of elevated EDPs due to accelerated ELN degradation or EDP-independent activation of NEU1 in the renal microvasculature^42,43^. Additionally, DANA potentiated the natriuretic and kaliuretic effects of furosemide in *Eln^+/+^* mice and male but not female *Eln^+/-^*mice. These data suggest that ERC signaling modulates NKCC2 and ENaC activity, and this modulatory effect is altered or diminished, sex dependently, in *Eln* haploinsufficiency. ERC blockade with DANA also enhanced urinary sodium and potassium excretion in the presence of V2R activation in *Eln^+/-^* mice, suggesting a functional role of ERC signaling in the regulation of ENaC and NKCC2 activity by ELN and its derivative bioactive peptides.

Our study and data interpretation are not without limitations. For instance, the effects of acute ECFV expansion and the simultaneous pharmacological manipulations of V2R and ERC on blood pressure were not evaluated. This could have given more insights into the pathophysiologic mechanisms of systolic hypertension resulting from *Eln* haploinsufficiency. Furthermore, the ERC blocker was administered systemically, as such extra-renal effects may have confounded the interpretation of the results. Future studies should aim at teasing out the molecular mechanisms by which signaling downstream of V2R and ERC intersect in the renal epithelium of various nephron segments to modulate the regulation of water and electrolyte homeostasis by the kidney. Together, the observations made in this study lead to the conclusion that *Eln* haploinsufficiency alters the functional link between signaling downstream of V2R and ERC for finetuning renal water and electrolyte handling, thereby leading to augmented sodium reabsorption and expanded ECFV (Figure 7). In future experiments, it would be interesting to test the potential therapeutic implication of our findings by assessing the anti-hypertensive efficacy of combining loop diuretics and ERC inhibitors compared to more commonly prescribed anti-hypertensives for age-related hypertension, including thiazides and calcium channel blockers.

**Figure 7.**
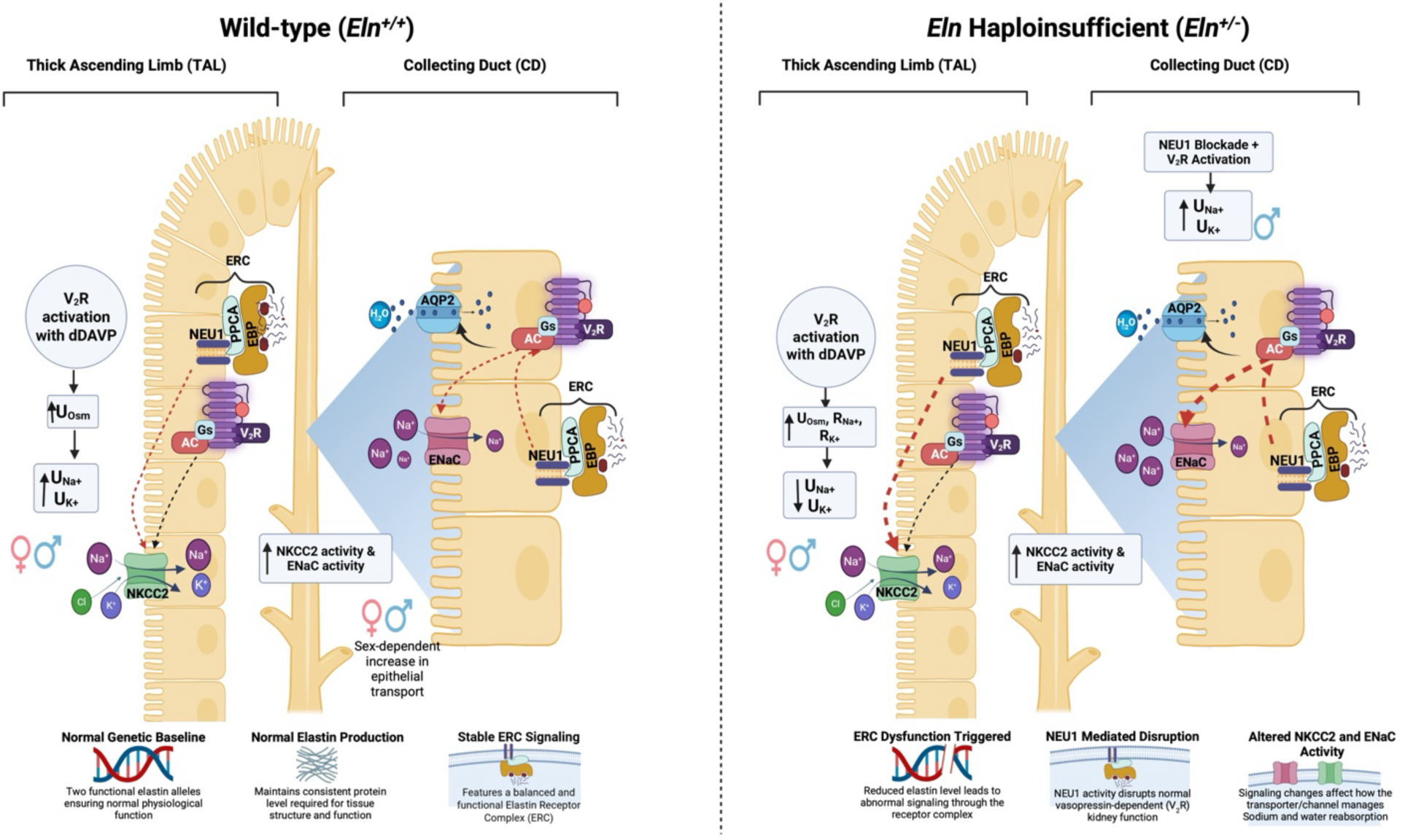
Putative mechanism of neuraminidase 1 (NEU1)-mediated modulation of vasopressin receptor-activated water and electrolyte handling by the kidney in *Eln* haploinsufficiency. AC, Adenylyl Cyclase; AQP2, Aquaporin 2; dDAVP, 1-deamino-8-D-arginine vasopressin/Desmopressin; EBP, Elastin Binding Protein; ENaC, Epithelial Sodium Channel; ERC, Elastin Receptor Complex; H_2_O, Water; K^+^, Potassium Ion; Na^+^, Sodium Ion; NKCC2, Sodium-Potassium-Chloride Cotransporter 2; PPCA, Protective Protein/Cathepsin A; R_K+,_ Reabsorbed potassium; R_Na+_, Reabsorbed sodium; U_K+_, Urinary potassium excretion; U_Na+_, Urinary sodium excretion; U_Osm_, Urinary osmolality; V_2_R, Vasopressin receptor 2.

## ACKNOWLEDGEMENTS

The authors thank members of the Osei-Owusu lab for technical assistance and Dr. Jeffrey R. Schelling for his constructive critiques of the manuscript.

## AUTHOR CONTRIBUTIONS

G.K., A.S.B., K.M, J.A.M., and P.O. performed experiments, and processed and analyzed data; G.K. and P.O. drafted the manuscript; G.K., J.A.M. and P.O. edited and revised the manuscript; all authors approved the final version of the manuscript.

